# Bees support farming. Does farming support bees?

**DOI:** 10.1101/804856

**Authors:** Preeti S. Virkar, Ekta Siddhu, V. P. Uniyal

**Author notes:** This author contributed equally to the work. This author also contributed equally to the work.

## Abstract

Tropical regions are subjected to rapid land use changes altering species composition and diversity in communities. The non-*Apis* bees are vital invertebrates continued to be highly neglected in the tropics. We compared their diversity status, richness and composition across natural areas and agroecosystems in Doon valley, a subtropical-temperate landscape situated at the foothills of outer Himalayas in India. We investigated how two major habitats relate to non-*Apis* bee diversity, specifically seeking answers to (1) Whether natural habitat is a refuge to richer and rarer bee communities than agroecosystems? (2) Are natural habitats important for supporting wild bee populations in agroecosystems? (3) Do polyculture farms behave similar to natural habitats and therefore support richer bee communities than monoculture? Observation and pantrap sampling were used to collect data. We recorded 43 species belonging to bees of five families. The findings of our investigation demonstrate the importance of natural habitats as a potential refuge for non-*Apis* bees. The findings highlighted that Doon valley harboured twenty-five rare species of non-*Apis* bees, and natural habitats are a refuge to 11 rare specialist species (clamtest; Specialization threshold K = 2/3, Alpha level = 0.005). Natural habitat diversity in Doon valley supports bee communities in nearby agroecosystems (R2 = 0.782, SE = 0.148, P = 0.004). Polyculture practices in agroecosystems (<100m from forest H’ = 2.15; >100m from forest = 2.08) in the valley mimic natural habitats (H’ = 2.37) and support diverse non-*Apis* bee communities (2.08) in comparison to monocultures (<100m from forest H’ = 2.13; >100m from forest =1.56). Bees evolved with flowering plants over 120 million years and they suffice an ever-growing anthropogenic nutrition needs with their services through enhanced agricultural production in pursuit of forage. We finally recommend similar assessments of bee diversity and plants they support in different habitats and vice versa.

## Introduction

Pollinators have a crucial role to play in pollination, a key ecological service, that enhances plant production [1, 2]. Bees, in particular, are considered the prime pollinator group [3, 4]. Beekeeping has hence become an indispensable part of farming cultures worldwide. Managed honey bees successfully pollinate and are responsible for the production of seeds and fruits of about 75% of the most commonly consumed food crops worldwide [5] and an unknown number of wild plants globally [1,2,5]. In addition to the managed honey bees, pollination by the native wild bees is an indiscernible but imperative process to the ecosystem. Native wild bees (non-*Apis*) other than honey bees (*Apis*) are increasingly finding the center stage in the backdrop of the global decline of the managed bees [6, 7]. Over the last half a decade the honey bee populations suffered a massive decline globally, predominantly in North America [8, 9] and a Europe [10, 11], leading to pollinator crisis [12].

There are several reasons for the bee decline such as climate change [13, 14], pollution, pesticide usage [15, 16], introduction of exotic species [17, 18], electromagnetic waves from mobile towers [19], and pathogens, [20] etc. Changing climate and land use are among the prime drivers of pollinator decline degrading their habitats. Conversion of additional natural land for agriculture is a significant cause of pollinator habitat degradation [21]. It alters biodiversity [22, 23], and is predicted to be one of the leading causes of species loss in the future owing to anthropogenic modifications in the global environment [24–26], particularly in the tropics. Tropical regions sustain rare and endemic species of plants, animals and their ecosystem services. Owing to lack of exploration and documentation, the discovery of many species may be yet remaining. There is limited baseline information on the status of non-*Apis* bees in the changing tropical and subtropical landscapes [27].

The global human population is estimated to reach nine billion by 2050 [28, 29]. Catering to the nutritional needs of a rapidly growing population will mean the conversion of natural habitats to agriculture [30]. Over the past twenty-five years (from 1990-2015) 129 million ha of natural land were lost to agriculture worldwide [31]. Tropical and developing countries such as India have undergone much rapid land cover conversions. From 1880-2010 the land use land cover (LULC) in India the forests cover decreased by 34% (134 million ha to 88 million ha) while the agricultural land increased by 52% (92 million ha to 140 million ha) [32]. Studies demonstrate that natural habitats are reservoirs of resources required by bees that may be absent in agroecosystems [7, 33–36]. Forest cover reduction and farmlands gaining vegetation uniformity with increasing monoculture cause the loss of nesting and foraging sites of the bee pollinators [37, 38]. Studies on semi-natural habitats have shown to differentially support bee populations in agricultural landscapes [39, 40]. Pollinators’ utilize forest and agrarian habitats for resources such as forage and nesting [22, 41–44]. Monoculture reduces not only natural areas [45] in and around farms but also the floral diversity [38]. The recent bee falloff is attributed to increased conventional monoculture agricultural practices that reduce wildflower abundance [16,46–49]. These practices upset the plant-pollinator community structure [14] and function [50]. Agricultural intensification is an critical global change presently leading to local and global consequences such as poor biodiversity, soil depletion, water pollution and eutrophication and atmospheric components [51]. It can negatively affect the insect pollination services, and thus, decreasing the production of 66% crops impacting the global nutrition supply [52]. In the wake of such information, one wonders what kind of differences agriculture and natural habitat might bring to bee diversity.

The Himalayan ecosystem is one of the global biodiversity hotspots. It is gradually falling prey to alterations in climate and land cover, to support growing economic needs. These very causes are affecting bee populations worldwide as well, and the Himalayas are particularly sensitive to them [53–56]. Rising temperatures influence plant reproduction and phenology. Mismatch of the plant life cycle will, in turn, affect the pollinators [13,57–59]. Understanding the ecosystem services in the Himalayas has begun [60]. The Global Pollination Project, in particular, investigated the contribution of insect pollinators to the productivity of important crops such as oilseeds, fruits and spices cultivated in the Indian Himalayas [61].

Himalayan agriculture is dependent on the managed pollinators for crop productivity. A secondary economic source through honey production is an age-old practice in these mountains [62–65]. The introduction of the non-native honey bees, increasing anthropogenic pressures on the natural resources, climate and lifestyle alterations are further challenging the traditional beekeeping in the region. Few investigations revealed honey bee declines reported in the Himalayan ecosystems [62, 65–68]. The rate of honey bee decline in the region and South Asia on a broader scale is unknown. This gap points out the need for utilizing the services of non-*Apis* bees to improve the reproduction in the wild and cultivated Himalayan plants. Lack of comprehensive investigation and historical records on non-*Apis* bees is a hindrance towards understanding their ecological function and economic potential in the region. Doon valley is a mosaic of natural, agricultural and human habitats, situated at the foothills of the outer Himalayas. Demonstrating both, traditional chemical-free temperate mountain farming and the conventional intensive cultivation from the Gangetic plains, this Himalayan valley is a perfect example of the changing Himalayan LULC. Few studies in the recent past that have looked at the insect pollinators in Doon valley. Jiju et al [69] recorded pollinators and flower visitors *viz.* Dipterans (n= 7), Hymenopterans (honey bees n=3-, wasp n=1-), Coleopterans (n=3), Hemipterans (n=1) and Lepidopterans (n=4) in an organic farm. This investigation, however, lacked any records of non-*Apis* bees. Migrating human populations from the mountains and the plains are causing rapid land use alterations affecting diverse faunal species in the Doon valley (Yang et al. 2013). Transitionary climatic conditions and geography make the Doon valley a unique landscape ideal for supporting a potential source population of non-*Apis* bees for the surrounding areas. We investigate how two major habitats of LULC viz. natural and agriculture in the Doon valley landscape, affect non-*Apis* bee diversity, specifically seeking answers to (1) Whether natural habitat is a refuge to diverse and rarer bee communities than the agroecosystems? (2) Are natural habitats important for supporting wild bee populations in agroecosystems? (3) Do polyculture farms behave similarly to natural habitats, thus support species-rich and diverse bee communities than monoculture?

## Methods

### Study Area

Doon valley (latitudes 29 °59’ to 30 °30’ N and longitudes 77 °35’ to 78 °24’ E) is situated in the western corner of Uttarakhand state, India (Fig 1). It is spread over approximately 1850 km2 area with the elevation ranging from 330 m to 800 m above mean sea-level. The valley is a mosaic of natural habitats such as forest patches, seasonal and perennial riverine systems, agriculture land and urban settlements (Fig 2 and Table 1). Situated at the foothills of the Himalayas, the valley is sandwiched between the Shivalik ranges. The rivers Ganga and Yamuna mark the south-eastern and north-western boundaries of the valley, respectively. Doon valley is a fragile, tectonically active landscape with a climate ranging from sub-tropical to temperate type. Ecologically, the valley forms a landscape-level ecotone between the hot tropical plains and the temperate Himalayan mountain ecosystems. Traditionally, the farming practice consists of multiple crops in different seasons of cultivation *viz.* Kharif, Jayad and Rabi. The major groups of crops grown consist of cereals (wheat, paddy, maize, and millets), oilseeds (mustard, sesame, and linseed), vegetables (potato, tomato, brinjal, radish, cabbage, okra, pea, onion, capsicum, French bean, ginger, garlic) and cash crops (sugarcane)[70]. The valley is famous for its variety of rice called Dehradun Basmati [71]and fruit orchards of litchi [72]. Other fruit orchards commonly cultured in the valley are pear, mango, guava, peach and Indian gooseberry. Numerous local citrus varieties and vegetables are grown seasonally in the valley for household consumption [70]. Based on the increasing demands and economic benefits, crops such as wheat and sugarcane are taking over the diverse cropping system in the Himalayas [73]. One can find many monocultures and polycultures in the valley based on the type and scale of farming.

**Fig 1.**
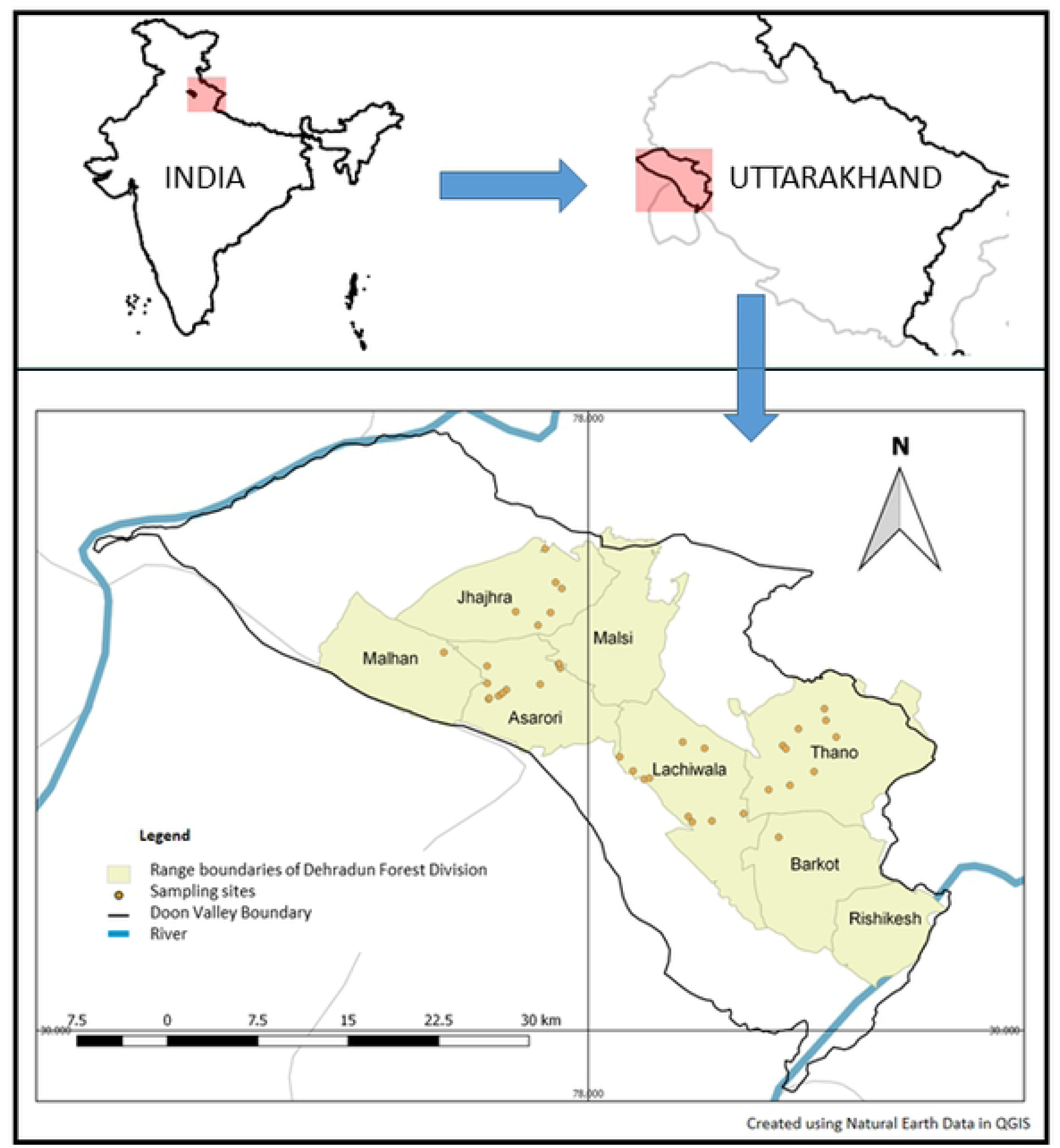
Sampling locations for bee species in Doon valley, India.

**Fig 2.**
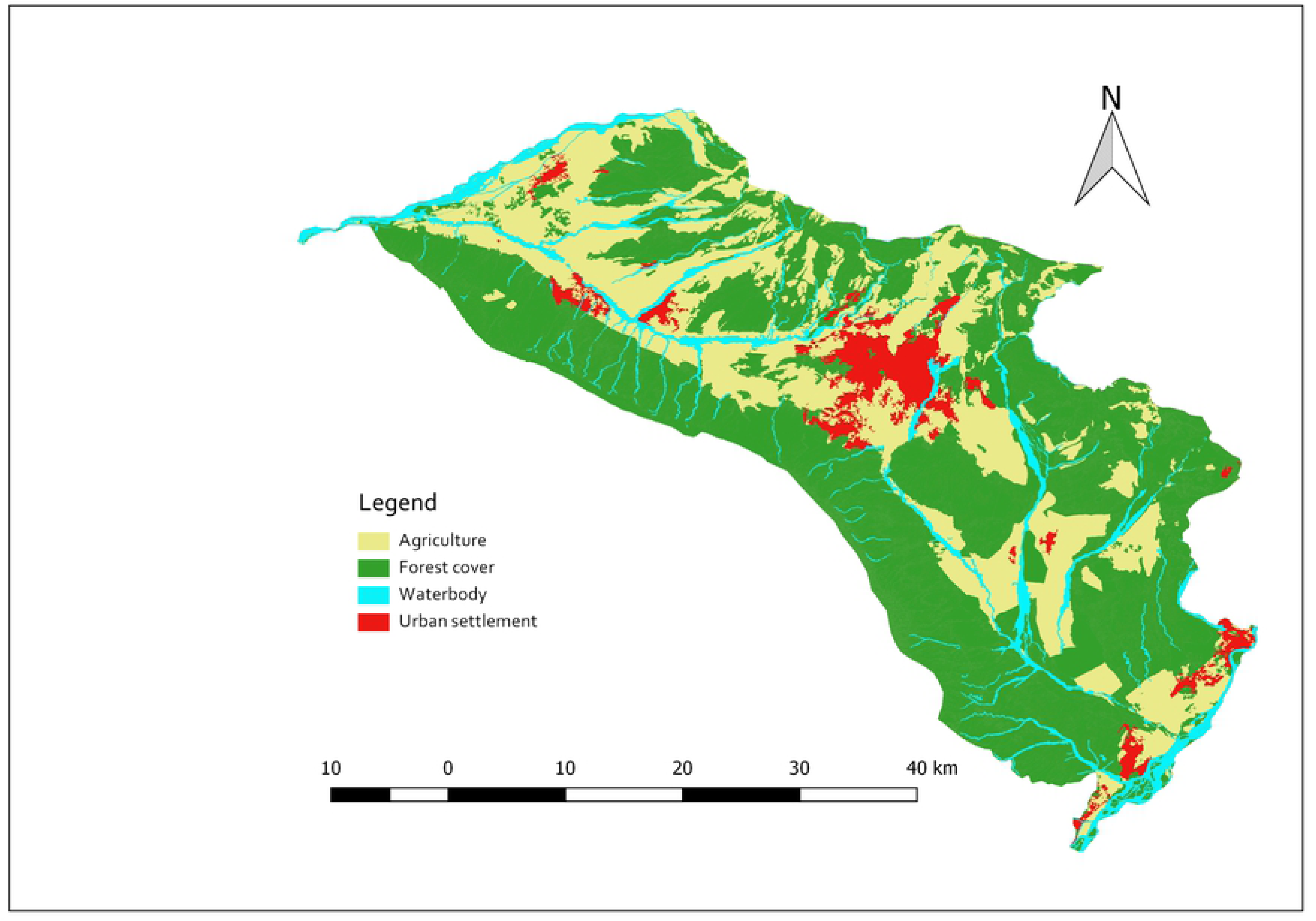
Land use and land cover map of Doon valley, India.

**Table 1.** Land use and Land cover (LULC) statistics of Doon Valley (in sq. km) in 2015 (Source: Mishra et al., 2015).

| LULC Class | Area in % | Area in sq. km |
| --- | --- | --- |
| Urban | 39.25 | 1205.37 |
| Water | 1.69 | 52.01 |
| Vegetation | 8.90 | 273.34 |
| **Agriculture** | 33.12 | 1016.98 |
| **Barren Land** | 17.04 | 523.29 |

### Study Design

Since we wanted to compare the bee diversity in natural and agriculture areas, we sampled majorly different types of the habitats mentioned above in Doon valley. The stratified random sampling design was followed in the two habitats (natural- 13500 sq. m and agriculture- 7000 sq. m) across the study site. The minimum radial distance of 1km was maintained between sites within a particular stratum. Sites were chosen such that a 50m buffer was left from the edge of the strata. Among the natural habitats, we sampled sal *Shorea robusta* forests, riverine patches and small patches of diverse vegetation (teak *Techtona grandis*, pine *Pinus roxburghii*, bamboo *Bambusa* sp. and derelict tea *Camellia sinensis* plantations). Polycultures and monocultures consisting of food crops and fruit orchards were sampled among the agricultural habitats. We laid 27 sampling plots in the natural habitat and 14 plots in the agriculture. The plots were belt transects of 100m×5m with five subsampling plots of 5m×5m dimension. The study site consisted of 203 subsampling plots in the natural (Sal- 45, riverine- 60 and miscellaneous-30) and agricultural (polyculture- 33, monoculture- 35) habitats. The sampling was carried out in the peak flowering periods. Three replicates were repeated from February to May in 2012, January to May 2013 and 2014.

Bee Sampling and Identification- Bees were sampled using active and passive methods on each transect. The active methods included visual observations using a transect walk and net sweeping. Visual observations were performed with a walk along the length of the transect searching for bees up to 2m on either side. Any sightings beyond this distance were ignored. It took approximately 20 minutes to complete one transect walk. Three temporal replicates of the transect walk were taken from 600 to 800 hrs., 1200 to 1400 hrs. and 1600 to 1800 hrs. Species unidentifiable in the field were collected using insect nets for further identification. The passive method comprised of the use of coloured pantraps to collect bees that are attracted to flower colours [74–77]. A set of three colours of plastic pantraps were used: yellow, blue and white [39]. Initially, the pantraps were left for 24 hrs. However, this duration yielded a low number to no bees. Thus, we left these pantraps for 48 hours and collected them on the third day. In total, 1827 pantrap sampling sessions were run (203 traps of three colours with three replicates) throughout the study period across the valley. Bees collected using all the different methods was appropriately curated. Specimens were stored either as a dried specimen and spread in insect boxes or as wet specimen in 70% ethanol vials till further identification. A standard identification key by Michener [4] was referred, to identify bees into families and further. We sorted the specimen into morphospecies as recognizable taxonomic units (RTU) for further data analysis [78–80].

### Data Analysis

Active sampling resulted in 2799 (Sweep Net=34, Observation=2765) bee records compared to passive (n=432) methods. We pooled the data obtained from both active and passive sampling to assess the species diversity of bees across different families. We calculated the bee species richness for all the habitats using non-parametric estimators (Colwell and Coddington 1994). We computed the Bray-Curtis dissimilarity index to examine whether bee species composition in the two primary habitats varied in Doon valley using Analysis of Similarity (ANOSIM) [81, 82]. A non-metric multidimensional scaling (NMDS) ordination was performed on the rank orders of dissimilarity values obtained from the ANOSIM. We encountered honey bees frequently (n = 2468) compared to non-*Apis* bees (n = 331) in active sampling, suggesting an observation bias towards the former. Hence, we used only pantrap records of bees for the ANOSIM. We performed a multinomial species classification based on species habitat preferences to test the association of bee compositions with natural and agricultural habitats [83]. The data for all the agricultural sampling plots at varying distances from natural habitats was compiled to explore whether natural habitats act as a refuge for bee communities of the agroecosystems. We classified the distance of sampling plot in agriculture from the nearest natural habitats into seven classes. Each distance class was 100 m wide beginning at zero to 700 m. Individual Shannon diversity indices (H’) were computed for each of the eight distance classes. We used regression analysis on the Shannon diversity for each distance class to investigate if nearness to natural habitat affects the diversity of bee assemblages in agroecosystems.

We wanted to test whether polycultures behave similarly to natural habitats compared to monocultures. For this we measured the Shannon-Weiner diversity of bees across 78 subsampling plots from monoculture and polyculture farms near (< 100m) and far from forests or wilderness (> 100m) *viz.* 20 monocultures near to the forest, 20 monocultures far from the forest, 18 polycultures close to forest and 20 polycultures far from forest. We compared the farm Shanon-Weiner diversity values with that of the different forests combined.

The analyses were carried out in the program R version 3.3.1 using package “vegan”[84, 85].

## Results

We recorded 3231 (432 in pantraps, 2799 through observation samplings) individual adult female bees. These individuals belonged to 43 species of bees falling in five families and 17 genera (Table 2). There were thirty-nine non-*Apis* and four *Apis* (*Apis indica, A. florea, A. mellifera* and *A. dorsata*) bee species.

**Table 2.** List of bees from Doon Valley.

| No. | Family | Bee Species | Agriculture<br>habitat | Natural<br>habitat |
| --- | --- | --- | --- | --- |
| 1 | Andrenidae | *Andrena Euandrena* sp. 3 | + | + |
| 2 | Andrenidae | *Andrena Euandrena* sp. 4 | + | + |
| 3 | Andrenidae | *Andrena flavipes* | + | + |
| 4 | Andrenidae | *Andrena* sp. 2 | + | + |
| 5 | Andrenidae | *Andrena* sp. 3 | + | - |
| 6 | Andrenidae | *Andrena Zonandrena* sp. 1 | + | + |
| 7 | Andrenidae | *Andrena* sp. 4 | + | + |
| 8 | Andrenidae | *Andrena* sp. 1 | + | + |
| 9 | Apidae | *Amegilla zonata* | - | + |
| 10 | Apidae | *Anthophora* sp. 2 | - | + |
| 11 | Apidae | *Anthophora* sp. 1 | - | + |
| 12 | Apidae | *Apid* sp. 1 | - | + |
| 13 | Apidae | *Apis dorsata* | + | + |
| 14 | Apidae | *Apis florea* | + | - |
| 15 | Apidae | *Apis indica* | + | - |
| 16 | Apidae | *Apis mellifera* | + | + |
| 17 | Apidae | *Bombus haemorrhoidalis* | + | + |
| 18 | Apidae | *Ceratina smaragdula* | + | + |
| 19 | Apidae | *Ceratina* sp. 5 | - | + |
| 20 | Apidae | *Ceratina* sp. 2 | - | + |
| 21 | Apidae | *Ceratina* sp. 3 | + | + |
| 22 | Apidae | *Ceratina* sp. 4 | - | + |
| 23 | Apidae | *Colletid* sp. 1 | + | + |
| 24 | Apidae | *Tetragonula* sp. 2 | + | + |
| 25 | Apidae | *Tetragonula* sp. 1 | - | + |
| 26 | Apidae | *Xylocopa astuens* | + | + |
| 27 | Apidae | *Xylocopa* sp. 1 | + | + |
| 28 | Halitidae | *Halictid* sp. 4 | - | + |
| 29 | Halitidae | *Halictid* sp. 1 | + | + |
| 30 | Halitidae | *Halictus* sp. 2 | + | + |
| 31 | Halitidae | *Halictus* sp. 3 | + | + |
| 32 | Halitidae | *Lasioglossum* sp. 2 | + | + |
| 33 | Halitidae | *Lasioglossum* sp. 1 | - | + |
| 34 | Halitidae | *Nomia interstitialis* | + | + |
| 35 | Halitidae | *Nomia* sp. 1 | + | + |
| 36 | Halitidae | *Nomia wetwoodi* | + | - |
| 37 | Halitidae | *Patellapis* sp. 1 | - | + |
| 38 | Halitidae | *Sphecodes* sp. 1 | + | + |
| 39 | Megachilidae | *Megachile lanata* | - | + |
| 40 | Megachilidae | *Megachile* sp. 1 | - | + |
| 41 | Megachilidae | *Megachile* sp. 9 | - | + |
| 42 | Megachilidae | *Megchile* sp. 8 | - | + |
| 43 | Megachilidae | *Osmia adae* | + | + |

The species composition of bees was significantly more similar within habitats than between habitats (R statistic = 0.20, p = 0.004) (Table 3). Bee composition in agroecosystems slightly overlapped with the different natural habitats. We pooled the data for the different natural habitats and overlaid it on that of agroecosystems. We found that arable lands shared the majority of the species with the different natural habitats in the Doon valley (Fig 3).

**Fig 3.**
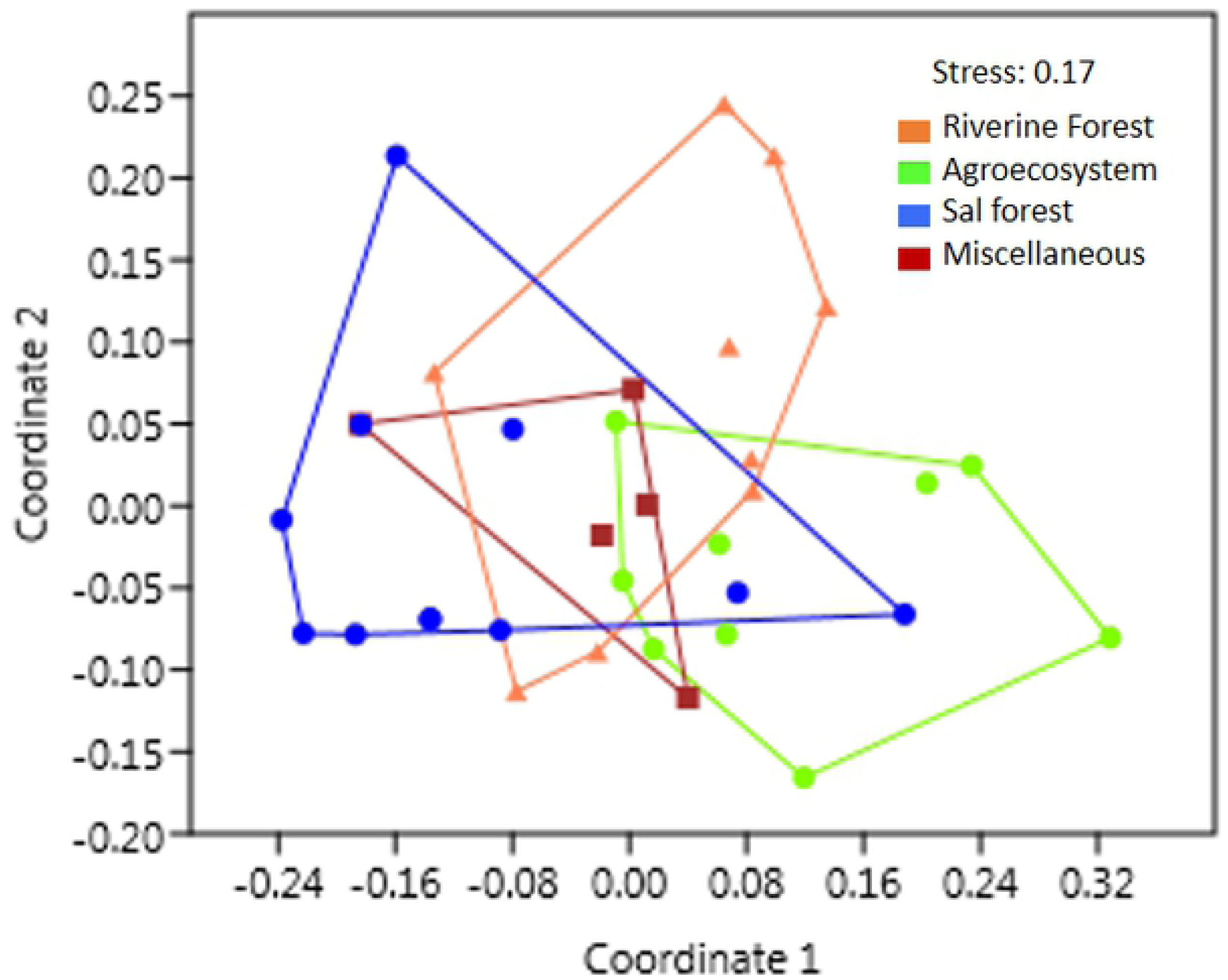
Species composition of bees in different habitats in Doon valley constructed using Nonmetric Multidimensional Scaling. Overlay of bee communities between agroecosystems (green polygon) and different types of forest habitats (blue, orange, & red). Forest habitats pooled together (black polygon).

**Table 3.**
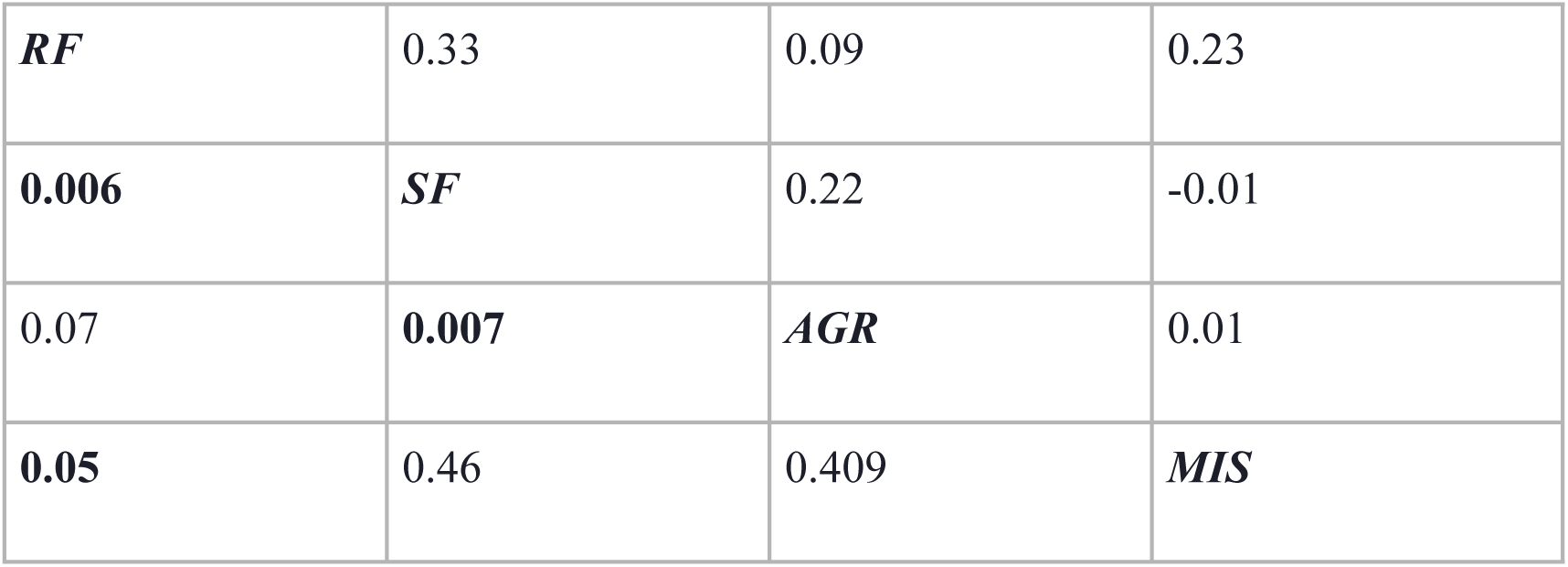
Comparison of bee community similarity between forest (Riverine Forest- RF, Sal Forest- SF and Miscellaneous vegetation- MIS) and agroecosystems (AGR) habitats in Doon valley (Pair wise ANOSIM, p<= 0.05). (Values above and below diagonal are R and p values respectively).

|  |  |  |  |
| --- | --- | --- | --- |
| ***RF*** | 0.33 | 0.09 | 0.23 |
| **0.006** | ***SF*** | 0.22 | -0.01 |
| 0.07 | **0.007** | ***AGR*** | 0.01 |
| **0.05** | 0.46 | 0.409 | ***MIS*** |

We classified the bees into habitat (agriculture, forest) specialists, generalists and species too rare to be classified with confidence (Fig 4). Of the total bee species, 58.14% were rare (n= 25), 25.58% (n= 11) were forest specialists, and 9.3% (n= 4) were generalists and found in both the habitats. During sampling three bee species (6.98%) *viz.*, Dwarf bee (*Apis florea*), Asiatic honey bee (*Apis indica*), and *Andrena* sp. three were detected only in the agricultural habitats. It is interesting to note that these species were observed inhabiting edges and interiors of the forests in the vicinity of the agroecosystems. Hence, these species are not specialists to agroecosystems but use both the habitats for foraging and nesting and may be subjective to availability of different resources year-round.

**Fig 4.**
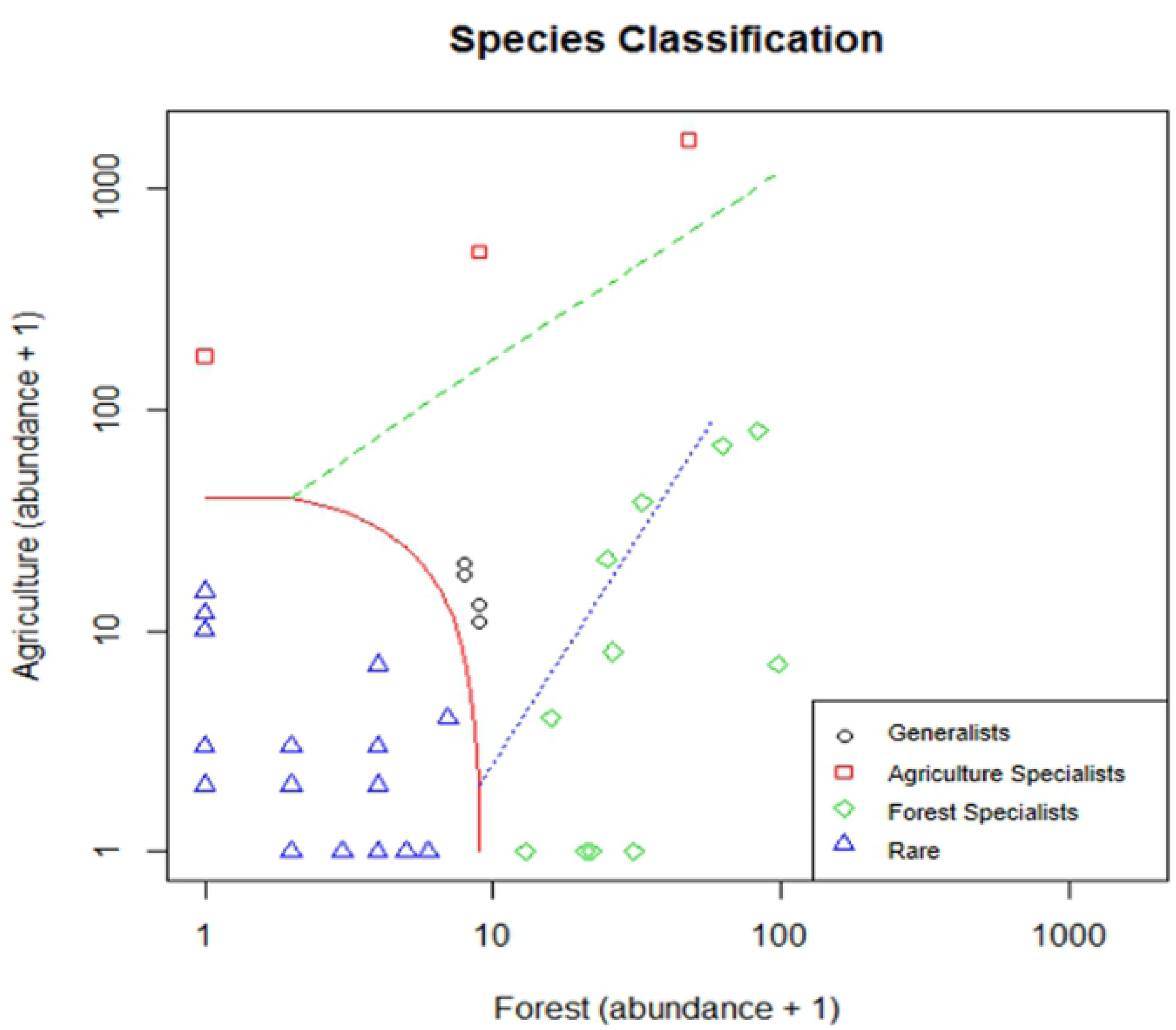
Species classification into specialists (agricultural and forest habitats) and generalists in Doon valley (Specialization threshold K = 2/3, Alpha level = 0.005).

Bee diversity was significantly higher in agroecosystems in close proximity to forests (H’ for <200 m = 1.60) compared to those further away (H’ for >600 m = 0.56) (R2= 0.782, SE= 0.148, p value= 0.004) (Fig 5).

**Fig 5.**
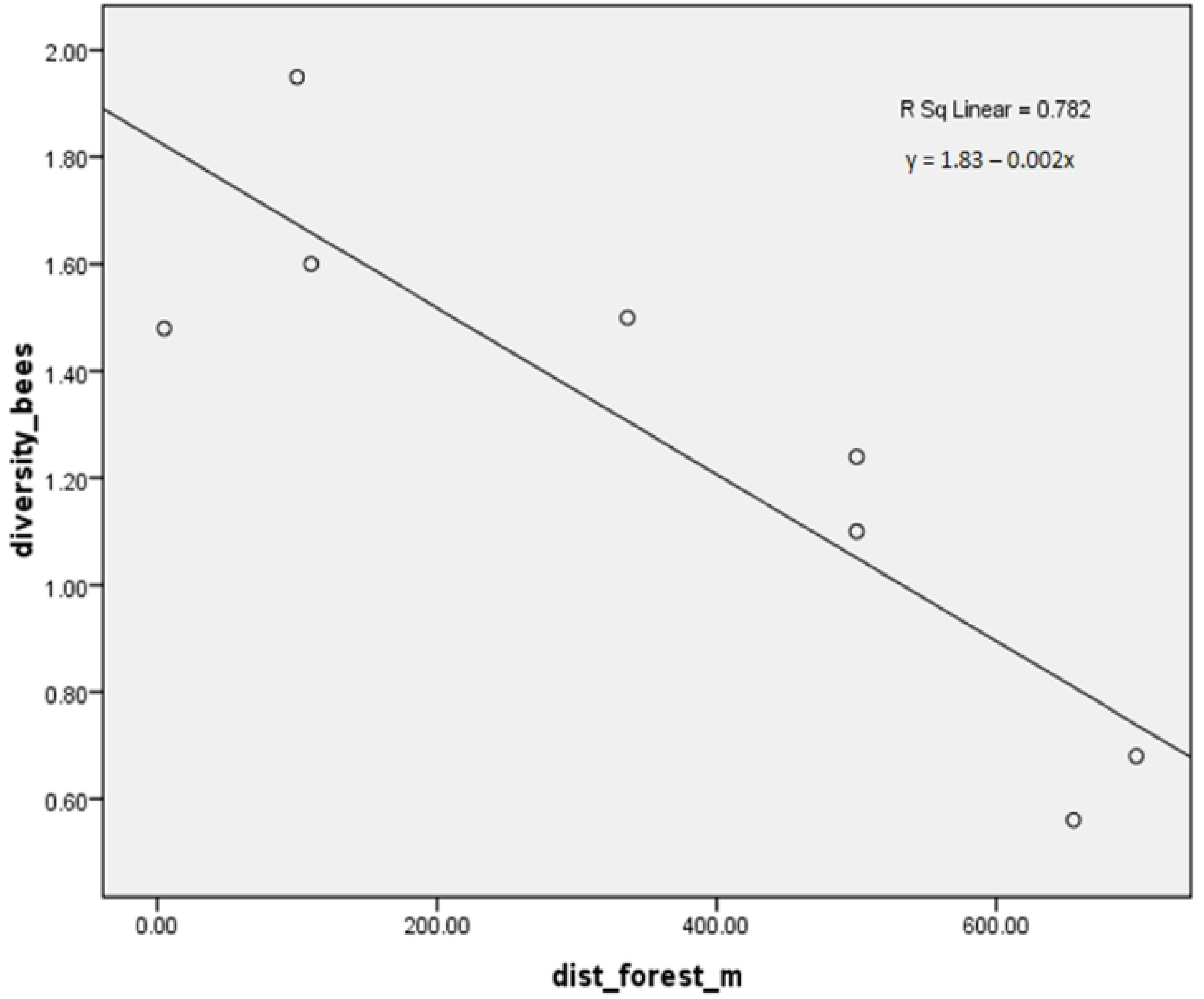
Influence of natural habitat on bee community richness in agroecosystems (linear regression model; R2 = 0.782, SE = 0.148, P = 0.004).

Our results demonstrate that forests harboured the highest number of bees (n=21), followed by polycultures near forests or wilderness (n=19) and monoculture near the forest (n=15), polyculture away from the forest (n=13) and monocultures away from the forests (n=9). The Shannon-Weiner diversity of forests was 2.37, monocultures near forests were 2.13, polycultures near forest was 2.15, monocultures away from forest were 1.55 and polycultures away from the forest was 2.08. The bee community diversity between monocultures and polycultures close to the forests were similar than those farther away. Polycultures supported a diverse bee community than monocultures that were away from the forests (Table 4).

**Table 4.** Shannon-Weiner diversity of bee communities in forests, monocultures and polycultures across Doon Valley, India.

| Habitat | Shannon-Weiner diversity | Species Richness |
| --- | --- | --- |
| Monoculture away from forest | 1.55 | 9 |
| Monoculture near forest | 2.13 | 15 |
| Polyculture away from forest | 2.08 | 13 |
| Polyculture near forest | 2.15 | 19 |
| Forests | 2.37 | 21 |

## Discussions

Continual land use changes to meet the needs of a growing human population are predicted to be the primary drivers of species decline [24], including essential pollinators such as bees [21]. The status of the vast majority of non-*Apis* bees remains uncertain owing to inadequate exploration. Honey bees, on the contrary, form only a small portion and yet have gained more considerable attention and became exhaustively studied. We attempt to bridge this gap and investigate the role of changing land use in shaping the non-*Apis* bee communities in a fragile river valley landscape at the foothills of the Himalaya. Ours is the first study on bees of this region and may be used as baseline evidence to compare future patterns in their communities locally. We found that Doon valley harbours 6.3% of the 678 bee species recorded from the Indian sub-continent [86]. Our investigation pointed out that a majority of these species consist of the less explored non-*Apis* bees (90.7%). Various bee researchers have demonstrated that honey bees (genera *Apis*) are a tiny fraction of the vast majority of bees described from the world over [4, 87]. It is not surprising that the Doon valley demonstrates a similar pattern of the bee biodiversity. Many (n = 30) bees recorded in the present study (n = 43) are segregated into morphospecies and are yet to be identified up to species level. Using specific taxonomic keys for bees may increase the total number of species in India.

We found that bee composition within habitats demonstrated more significant similarity than between habitats in Doon valley. Interestingly, the bee composition of the agroecosystems was a subset of the natural habitats. Bees require a diverse habitat with resources for breeding, nesting, and foraging to complete their life cycles. All the resources required may be unavailable within the habitats that bees occupy. For example, some bees depend on particular plant pollen for proteins that may be highly nutritious for reproduction [88]. However, flowers with such protein-rich pollen may not bloom year around always, which makes them move from one habitat patch to another. Agroecosystems are abundant sources of mass flowering and may serve as a food support system for bees at a particular time of the year. Thus, numerous bees utilize both natural and human-managed ecosystems such as cropland to suffice their nutritional, sheltering and their reproduction requirements [22,41,43,44,88]. We sampled the bees in the peak flowering period in Doon valley. During this is the period many farms in the valley have abundant vegetables, fruits and oilseed plants in the various stages of flowering. Although most bees nested in different natural habitats, agroecosystems may act as rich foraging grounds for some species during the spring in the valley. Thus, the bee composition of agroecosystems overlapped partially with that of the natural habitats in the valley.

Our study highlighted that more than half of the bees in the valley were rare. Patterns of the rarity of bees through a large number of singletons reported may arise because of under-sampling, an unrecognized universal trait of a rarity in communities or existence of transient species that thrive well in other regions [89]. Williams et al. [89] tried to understand the change in bee communities across the globe. They highlighted that the native bee community are highly diverse and rich in rare species at local scales (due to a large number of singletons at such scales). Several other researchers reported this pattern in various studies across regions [90–94]. We sampled the valley in peak flowering season, which may answer a part of the question of a rarity in species due to under-sampling during other seasons. Considering that Doon Valley is a transition between two different ecosystems, the other two possibilities of a large number of a rarity in communities based on a universal pattern and transient species cannot be ruled out.

Over one-fourth of the bees, we recorded from the valley preferred natural habitats of the forests. Few generalist species were found in both the habitats. Interestingly, 3 of the bee species *viz.* Dwarf bee (*Apis florea*), Asiatic honey bee (*Apis indica*), and *Andrena* sp. 3) were classified to prefer agroecosystems. *Apis florea* prefers bushy shrubs with strong twigs to make their nests. Thus, they are highly sensitive to incidences, such as fires. Forest fires are frequent in the valley from mid to end of the spring. Activities such as leftover cigarette buds discarded by humans venturing into the forests for extracting fodder and firewood usually trigger forest fires. They are intensified further by the dry season. Hence, farm boundaries and hedgerows form favourable spaces for the dwarf bees to build their nests beside the availability of abundant food in the spring. The Asiatic honey bees are generalist feeders and feed in large numbers in farmlands with abundant flowering crops. One can observe these bees foraging in large numbers on mustard and other flowering crops in Doon valley in the spring. During different seasons these bees are seen foraging on a diversity of wild and cultivated plants. They reside in parallel combs similar to their close cousins, the *Apis mellifera* in hollows of trees, rocks or undisturbed enclosed spaces in case of urban areas. Finding *A. indica* hives in farmlands are rare unless there are intact patches of large trees with hollows or rocky patches. Forests are suitable and typical habitats to find these bees. *Andrena* bees nest on well-drained, steep open banks [95]. Field boundaries that are used to segregate different crops in Doon valley provide such habitats in abundance. Morandin et al. [40] reported similar observations in their study where untilled field margins adjoining roads provided suitable habitats for *Andrena* bees to nest. Hence, these three species should be treated as facultative rather than specialists as they utilize resources from both the forests and the agroecosystems.

Agroecosystems at increasing distances from the natural habitats demonstrated lower bee diversity in our study. Numerous studies in the past reported similar findings. Pollinators depend on natural and semi-natural habitats adjacent to agroecosystems for supplemental food resources and shelter [6,96–98]. Wild bees in forests demonstrate limited foraging ranges. The valuable services of these bees influence agroecosystems in the vicinity of forests [7]. Maintaining natural habitats within or surrounding the agroecosystems crucial for wild bee pollination services [99].

Our results illustrate that monocultures close to forest ecosystems show diverse bee communities than those farther away. Surprisingly, the bee richness in polycultures away from the forests was similar to monocultures in close vicinity of the forests. Our results demonstrate that polycultures behave similar to natural habitat in conserving bees. The outcomes of our study indicate that polyculture farms had rich bee species compared to monoculture. Earlier studies (Potts et al. 2003) showed that bee communities are positively linked to floral diversities and prefer polyculture over monoculture [100]. The traditional Himalayan agriculture consists of diverse crops [101]. Polyculture systems that we sampled consisted of different crops in combinations of 2 to 4 species and wildflowers on the bunds to segregate them, which may have supported a rich composition of bees. On the contrary, our study recorded fewer bee species in monocultures of wheat (*Triticum* spp.) and mustard (*Brassica juncea* (L.) Czern). Wheat is wind-pollinated and does not provide biotic pollinating agents with rewards such as nectar and pollen. In monoculture, we found most bee species on mustard or on wild invasive plants such as *Lantana camara* and *Ageratum conyzoides* that grew on the boundaries of the farms. Polycultures have a greater diversity of native plants than monocultures which influences the bee biodiversity in agroecosystems [102]. Our findings demonstrate that forests are reservoirs of diverse bee communities in adjoining patches of agroecosystems. The present analysis supports the fact that polycultures act as reservoirs of habitat to bees compared to monocultures in close vicinity to natural habitats.

## Conclusions

Our investigation on bees in Doon valley, a landscape with gradual changing LULC at the foot of the Himalaya, is the first one to document the bees (*Apis* and non-*Apis*) in the forest and agricultural habitats of the region. Our findings demonstrate that the Doon valley landscape and its natural habitats are a refuge to a diverse community of rare and specialist bees. Agroecosystems shared a lot more species in common with each of the natural habitats. Forests and wilderness are essential habitats to support diverse bee communities in and around the agroecosystems. Furthermore, polycultures support higher bee diversities over monoculture practices. Wilderness along field boundaries such as hedges stands as connectivity between different habitats. Monoculture expansions have diminished these vital corridors [38,103,104], affecting the survival of pollinators [45]. Our study implies the importance of multi-crop farming and preserving as many natural habitats as possible to attract native bees and benefit both cultivated and wild plant production.

The global tree cover shows an increase of 7% (2.24 million sq. km) from 1982 to 2016 (Song et al. 2018), differing from a previous assessment claiming loss of forest (FAO 2015). India was ranked first in short vegetation gain (270,000 sq. km), most of which can be attributed to the agricultural intensification under the green revolution [105]. Extension of land under intensive agrarian practices is a prime cause of deforestation and loss of wilderness habitats. Wilderness play buffers to biodiversity that have lost intact natural habitats. Loss of wilderness causes the extinction of several terrestrial species across the global biogeographic realms, including the Indomalayan region [106]. Loss of wilderness habitats comprising of essential resources can drive local pollinators depending on them to extinction. Our investigation highlights that natural habitats are integral to bee diversity. In the milieu of honey bee decline worldwide Polycultures give some hope to non-*Apis* bee diversity in a Himalayan valley landscape. Thus, there is a ray of hope to “land sharing and sparing” as Baurdon and Giller to feed a growing population [107]. Using ecologically friendly farming such as polycultures nutritious biodiverse food for a growing population can be produced while conserving wild pollinators simultaneously. An assessment on similar lines is needed to understand bee diversity occupying natural and arable habitats in the tropical and sub-tropical regions of the world. Besides, it is crucial to understand what percentage of these bees support crop production and do our present agricultural practices in return support bees?

## Acknowledgements

We are grateful to the Director, Wildlife Institute of India, Director and Staff of Dehradun Forest Division, Uttarakhand. We thank the Department of Entomology, Indian Agricultural Research Institute, New Delhi for the guidance and identification of the specimen. We would like to extend our gratitude to Research Foundation for Science, Technology and Ecology, Navdanya Trust, New Delhi for their support to work with farmers. We are grateful to the State Biotechnology Department, Government of Uttarakhand, India. (SBD/R&D/01/11/02-87) for funding the study. We are indebted to the farmers in Doon valley for their cooperation in letting us conduct uninterrupted field work for the study.

